# Hide and seek in a rainforest canopy : when a fruit-eating carnivore (almost) met its top predator carnivore

**DOI:** 10.64898/2026.11.05.763601

**Authors:** Pierre-Michel Forget, Raphaëlle Abensur, Malika Gottstein, Coraline Pelon, Valentine Alt, Jocelyn Cazal, Lionel Picart, Eric Guilbert, Christophe Baltzinger

**Affiliations:** Muséum National d’Histoire Naturelle, UMR 7179 MECADEV CNRS-MNHN, LABEX DRIIHM, 43 rue Buffon, Paris, 75005, France; Arboriste, Roura, 97311, Guyane; INRAE, UMR ECOFOG, Kourou, 97310, Guyane; HEVEA, Malataverne, 26780, France; INRAE, EFNO, Domaine des Barres, Nogent-sur-Vernisson, 45290, France

**Keywords:** Camera trap, canopy, French Guiana, *Potos*, predator avoidance, prey hunting, *Puma*

## Abstract

Prey-predator interactions in the wild are spectacular. Common reported cases come from ground-dwelling animals, in open areas and by daylight. Here, we report, for the first time, a sequence with the nocturnal arboreal kinkajou (*Potos flavus*), being followed ca. 20 minutes apart by one of its predators, the puma (*Puma concolor*), 33 m aboveground in a rainforest in French Guiana.

## INTRODUCTION

Growing up with the story of Mowgli in The Jungle Book (Kipling 1894), as field biologists in tropical biomes, we often both dream and fear crossing the path of medium-to-large carnivores such as Bagheera and Shere Khan, the so-called big cats. These may either forage on the ground, hidden in the understory, or on a branch, resting in the low-stratum canopy, well positioned to encounter potential preys, human beings included (Garcia-Alaniz et al. 2010, Alldredge et al. 2019, Arias et al. 2024). However, not all large carnivores are able to climb trees, allowing medium-sized species such as the leopard (*Panthera pardus*) to “*drag prey into the upper branches of trees for storage*” (Vander Wall 1990) where it is protected from pilferage by larger cats. However, such interactions in closed habitats and high up in the trees, like in tropical forests, are rarely documented due to the difficulty of observations in space and time, particularly by night. The body mass of the leopard is within the same order of magnitude (17-65 Kg) as that of the puma (*Puma concolor*, 29-120 kg), but lower than that of the jaguar (*Panthera onca*, 31-158 kg) and tiger (*Panthera tigris*, 91-423 kg) (Emmons and Feer 1990, Etnyre et al. 2011). Therefore, not surprisingly, to avoid a deadly interaction with the tiger Shere Khan on the ground, «*Bagheera would lie out on a branch and call, “Come along, Little Brother,” and at first Mowgli would cling like the sloth, but afterward he would fling himself through the branches almost as boldly as the gray ape”* (Kipling 1894). Without doubt, being able to move high in treetops would allow humans and non-human primates, as well as many other small-to-medium sized canopy dwellers, escaping ground-dwelling large-sized predators. Nonetheless, tigers are known to be good tree climber, though may only venture a few meters above the ground to catch their prey (Nair 2024). Other large cats such as the leopard, puma and jaguar may also prey on primates (Chinchilla 1997, Corredor-Ospina et al. 2021, Dymit et al. 2026). There are casual evidences that leopards may hunt - not always with success - arboreal preys such as bonobo (*Pan paniscus*) on branches about 10–12 m high in a *Dialium* sp. tree in Central Africa (Corredor-Ospina et al. 2021). A tiger has also been observed unsuccessfully chasing a langur (posssibly *Semnopithecus hector*) on a small tree in India, a rare event given the known diet composition of tigers (Acharya et al. 2026). Questions thus arise about interactions between ground-dwelling top predators and their canopy-dwellers preys. Are the latter really safe from climbing large cats in the forest canopy? How high can top predators climb into the treetops?

**FIGURE 1:**
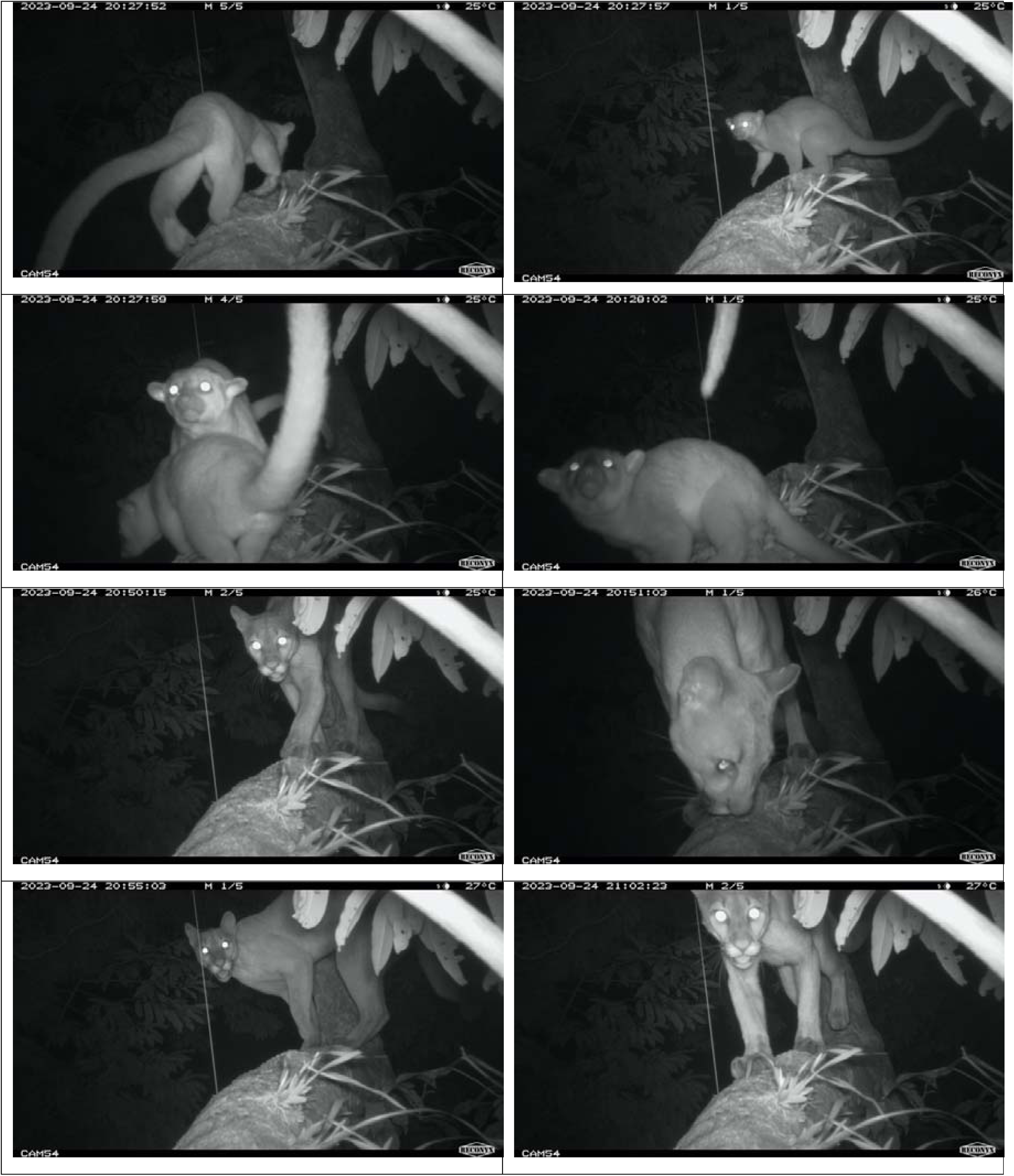
Kinkajous (*Potos flavus*) and Puma (*Puma concolor*) photographed with RECONYX® Hyperfire2 camera traps set at 33 m in height into the canopy of nutmeg tree (*Virola kwatae*, Myristicaceae) on the 24/09/2023 between 08:27:49 PM and 09:06:30 PM at Saut Pararé forest, Biological Station of Nouragues, National Natural Reserve of Nouragues, French Guiana. (Top two rows: adult female kinkajou, then joined by a juvenile. Bottom two rows: puma sniffing the spot previously visited by the kinkajous, and repeatedly visiting the spot at two closed instances).

Pumas standing or resting on lower branches or at the top of dead trunk in the temperate are a commonplace in image database such as Alamy (2026). As the number of visitors increases in the wild, there is a greater chance that someone may broadcast an interaction between a puma and its prey (e.g. Hedger 2022). In one daytime record, a puma attacked a sloth (*Bradypus tridactylus*) climbing a *Cecropia* sp., in a location not mentioned (National Geographic Wild 2002), a fast-growing tree with a small diameter and a small stature. Filming such an attack few meters above the ground is a very rare event (Guevada 2021), that is very unlikely to be documented in the tall tropical rainforest. Very rarely, a human observer may photograph a big cat high in the rainforest canopy, foraging, or even hunting their preferred preys (Dymit et al. 2026). In Kenya, a cheetah (*Acinonyx jubatus*) chasing and catching impalas (*Aepyceros melampus*) is a well-documented example (Veronesi 2007). Setting up camera traps in the canopy of tropical trees may overcome the spatial and temporal constraints associated with observing such interactions. Here, we report, for the first time, a sequence involving the arboreal kinkajou (*Potos flavus*), a nocturnal fruit-eating Carnivore, being followed by one of its predator, the puma (*Puma concolor*), 33 m aboveground in the crown of a nutmeg tree (*Virola kwatae*, Myristicaceae).

## STUDY SITE AND METHODS

Our observations took place in the tropical rainforest at the Saut Pararé forest within the Biological Field Station of Nouragues (Bongers et al. 2001) located 100 km south of Cayenne, the Prefecture of French Guiana. As part of the GUYACAM vertebrate long-term monitoring program in Guianan tropical forests (GUYACAM 2017-2026a), which aim to identify and quantify fruit-eating mammals and birds visiting fruiting nutmeg trees by coupling camera traps at the bottom and in the canopy of nutmeg trees, we equipped each selected trees (N = 10) with one camera trap on the ground and one in the canopy, most often with mature fruits (Abensur et al. 2026). After a rope was installed on the 19/01/2023 between 08:30 and 09:00 in a 41-m tall nutmeg tree (60-70 cm DBH), a RECONYX® HYPERFIRE 2 COVERT camera trap (serial number HLRXMQ07161697 (internally numbered GUYACAM #54), was fixed to the trunk by one of us (E. Guilbert) at a height of 33 m, facing a main horizontal branch (25 cm in diameter). On 11/06/2024, the camera trap was removed and replaced with another camera trap (same model, GUYACAM #67), by another author (C. Baltzinger). The animal species visiting the tree were later identified (by R. Abensur & C. Pelon) after the images had been stored at Agouti.eu (GUYACAM 2017-2026b). The camera trap remained operational and continued taking images until 08/04/2024, when the battery ran out.

## RESULTS

On 24/09/2023 at 20:27:49 a female adult kinkajou arrived, moved along the branch, stopped at the reiteration base, then abruptly returned to be joined within 10 seconds by a juvenile individual. Together, they rapidly moved upward, away from the lens, following each other within a couple of seconds at 20:28:00. Nearly twenty minutes later (20:49:43), one individual puma entered the scene, arriving from the crown periphery facing the camera trap, and remained in the vicinity of the branch during a quarter of an hour. At first, it spent ca. 30 seconds exploring the branch, especially sniffing the exact spots where the kinkajous had previously been recorded. It then disappeared at 20:51:36 to reappear at 20:54:16 in front of the camera trap, moved along the branch, turned back, and returned with the mouth open, to move away again at 20:55:35. It came back at 21:02:23, exploring further the branch with several roundabouts between the branch extremity and the trunk, apparently marking while passing closed to the tree reiteration, to finally leave at 21:06:30.

## DISCUSSION

Because large-bodied carnivores have large home ranges, studying their feeding ecology and habitat use requires appropriate methods and large-scale protocols to increase the probability of encountering the animals, or at best, the prey remains in their feces (Emmons 1987, Soria-Díaz et al. 2010, Ávila-Nájera et al. 2018, Craighead et al. 2022, Karandikar et al. 2022). In tropical America, however, the analysis of cats scats can be misleading because of the similarity in scat size when they co-occur (Emmons 1987). Nonetheless, a recent investigation using fecal DNA metabarcoding in Guatemala showed that jaguars are specialized on large terrestrial prey while pumas exhibit high consumption of arboreal primates (Dymit et al. 2026), a result consistent with previous study of cat scats in Corcovado National Park, Costa Rica (Chinchilla 1997). On Barro Colorado Island, Panama, Swinkels et al. (2022) used a network of camera-traps to monitor temporal proximity of ground-dwelling predator and prey animals. They found that the “*time until a prey animal appeared at a location was significantly longer than expected by chance if an ocelot had passed, and that the time until an ocelot appeared at a location was significantly shorter than expected by chance after prey passage*”. Documenting similar prey-predator interaction in the canopy would then be very challenging, but not impossible today thanks to safe tree climbing and arboreal camera trapping (Picart et al. 2014, Moore et al. 2021). During the course of the GUYACAM program, we had set fruiting trees with dozens of camera traps on the ground and in the canopy during 12-month cycles (Abensur et al. 2026). Although our protocol aimed at documenting the diversity and activity of fruit consumers and seed dispersers in the targeted fruiting trees (Coutant et al. 2022, Séguigne et al. 2022), it also favored describing the occurrence of other vertebrates species inhabiting the forest, without direct link to the focal study tree species. Overall, between 2017- 2026, we generated a total of 39,037 independent events (30’) at the targeted nutmeg trees (R. Abensur, pers. obs.), e.g. 24,237 events in the understorey, and 14,800 events in the canopy. Of these, we observed 33 margays (*Leopardus wiedii*), 145 ocelots (*Felis pardalis*), 21 jaguarundi (*Herpailurus yagouaroundi*), 41 pumas, and 33 jaguars. We observed only one interaction between a felid and its potential prey, namely the kinkajou-puma interaction, at a height of 33 m.

Although our protocol aimed at documenting the diversity and activity of fruit consumers and seed dispersers in the targeted fruiting trees (Coutant et al. 2022, Séguigne et al. 2022), it also favored describing the occurrence of other vertebrates species inhabiting the forest, without direct link to the focal study tree species. Indirect links may be the interaction between a fruit consumer vertebrate and its predators, and avoidance–attraction behavioral interactions as a whole (Swinkels et al. 2022, Dymit et al. 2025). In the canopy, we observed that kinkajous are abundant in tall *Virola* trees greater than 30 m (Séguigne et al. 2022, Abensur et al. 2026), and well adapted to foraging into the tree canopy (Koepfli et al. 2007) where they can supply themselves with fruit during the wet season (Julien-Laferrière 1993). The time until the puma appeared at our study *Virola* tree was very short (ca. 20’) after kinkajou passage which is consistent with prey tracking and predator avoidance (see Swinkels et al. 2022, Dymit et al. 2025). Kinkajou is adapted to a nocturnal arboreal life using a long prehensile tail allowing to forage high in the canopy where it shares *Virola* fruit resources with other diurnal arboreal consumers. In conclusion, our unique observation shows that kinkajou may also rapidly avoid its predator, thanks to prehensile tail-associated behaviors while moving upward into the tree crown (Mensah et al. 2024, Naas et al. 2025) that is supposedly less likely to be foraged by big cats.

## Supporting information

The entire sequence of images dated 24/09/2023 lasting from 08:27:49 PM and 09:06:30 PM.

## ACKNOWLEDGMENTS

We thank the Biodiversity French Office, the CNRS and the Labex DRIIHM, LEEISA Research Unit and the Biological Research Station of Nouragues, the UMR 7179 CNRS—MNHN, the INRAE, HEVEA Elagage, and the Fundation Crédit Mutuel Alliance Fédérale for their financial support of the GUYACAM project.

## Data Availability Statement

Data are presented in the MS and supplementary data

## Supplementary information

The entire sequence of images dated 24/09/2023 lasting from 08:27:49 PM and 09:06:30 PM.

